# Single-cell transcriptome analysis of fish immune cells provides insight into the evolution of vertebrate immunity

**DOI:** 10.1101/074344

**Authors:** Santiago J. Carmona, Sarah A. Teichmann, Lauren Ferreira, Iain C. Macaulay, Michael J.T. Stubbington, Ana Cvejic, David Gfeller

**Affiliations:** Ludwig Center for Cancer Research, University of Lausanne, Lausanne, Switzerland; Swiss Institute of Bioinformatics (SIB), 1015 Lausanne, Switzerland; European Molecular Biology Laboratory European Bioinformatics Institute, Wellcome Trust Genome Campus, Hinxton, Cambridge, UK; Wellcome Trust Sanger Institute, Wellcome Trust Genome Campus, Cambridge, UK; Department of Haematology, University of Cambridge, Cambridge, UK; Wellcome Trust – Medical Research Council Cambridge Stem Cell Institute, Cambridge, CB2 1QR, UK; Sanger Institute–EBI Single-Cell Genomics Centre, Wellcome Trust Genome Campus, Hinxton, Cambridge, UK

**Keywords:** single cell RNA-Seq, zebrafish immune cells, immune system evolution

## Abstract

The immune system of vertebrate species consists of many different cell types that have distinct functional roles and are subject to different evolutionary pressures. Here, we first analysed gene conservation of all major immune cell types in human and mouse. Our results revealed higher gene turnover and faster evolution of trans-membrane proteins in NK cells compared to other immune cell populations, and especially T cells, but similar conservation of nuclear and cytoplasmic protein coding genes. To validate these findings in a distant vertebrate species, we used single-cell RNA-Sequencing of *lck:GFP* cells in zebrafish to obtain the first transcriptome of specific immune cell types in a non-mammalian species. Unsupervised clustering and single-cell TCR locus reconstruction identified three cell populations, T-cells, a novel type of NK-like cells and a smaller population of myeloid-like cells. Differential expression analysis uncovered new immune cell specific genes, including novel immunoglobulin-like receptors, and neofunctionalization of recently duplicated paralogs. Evolutionary analyses confirmed a higher gene turnover and lower conservation of trans-membrane proteins in NK cells compared to T cells in fish species, suggesting that this is a general property of immune cell types across all vertebrates.

## Introduction

The immune system of vertebrate species has evolved into a highly complex structure, comprising many different subtypes of both innate and adaptive immune cells. Adaptive immune cells are broadly classified into B and T lymphocytes that can directly recognize and bind antigens with great specificity. Innate immune cells include a variety of myeloid cells such as monocytes, neutrophils, basophils, eosinophils and mast cells. A third major type of lymphocytes, the Natural Killer (NK) cells, has also been historically classified among innate immune cells (Sun and Lanier 2009; Sun et al. 2009). Traditionally, different immune cell types are distinguished based on unique combinations of cell surface markers. In mouse and human, many antibodies for these markers are available and can be used to isolate homogeneous immune cell populations using flow cytometry. Gene expression profiling studies of isolated immune cell populations have further allowed genome-wide identification of cell-type specific genes (Shay and Kang 2013; Watkins et al. 2009; Chambers et al. 2007; Vu Manh et al. 2014). These studies revealed an overall conservation of immune cells’ gene expression between mouse and human (Shay et al. 2013). However, beyond mouse and human, less is known about the characteristics and evolution of immune cell types mainly due to the challenges of isolating different immune cell populations.

Evolutionary studies based on mouse and human genes have shown that immune-related genes tend to evolve faster than other genes (Bailey et al. 2013; Boehm 2012; Flajnik and Kasahara 2010; Kosiol et al. 2008). This faster evolution may reflect a need of immune cells to adapt to a rapidly changing environment and specific pathogens. In addition, different immune cell types are subject to different evolutionary constrains. T and B lymphocytes can generate an extraordinary diverse repertoire of antigen-specific receptors as a consequence of *rag-*mediated somatic V(D)J rearrangement, and this process is conserved across all jawed vertebrates (Boehm 2012). Many orthologs of T cell specific genes, like CD4, CD8 and TCR genes, have been identified in all jawed vertebrates. In species like zebrafish, the V(D)J variable regions has been recently annotated (Meeker et al. 2010; Schorpp et al. 2006; Iwanami 2014). NK receptors instead are germline-encoded. Therefore, selection pressure to generate different receptor specificities and transduce signals is expected to operate at the population rather than at the individual cell level. Indeed, mammalian NK cell receptors have expanded and diversified in a species-specific fashion, like in the case of KIR receptors in primates and Ly49/Killer lectin-like receptors in rodents (Carrillo-Bustamante et al. 2016). NK-like cells have been identified in non-mammalian species such as chicken (Jansen et al. 2010), xenopus (Horton et al. 1996) and catfish where spontaneous killing of allogeneic cells by non-TCR expressing cytotoxic cells was demonstrated (Shen et al. 2004; Yoder 2004). A recent study using single-cell qPCR based on known markers of blood cell lineages revealed the presence of a small sub-population of immune cells in zebrafish, which were proposed to represent putative NK-like cells based on expression of NK-lysin genes (Moore et al. 2016). The identification of membrane receptors with similar genomic organization as the KIR genes in human provided additional evidence for the existence of NK cells in fish species. In zebrafish, these receptors include *nitr* (Yoder et al. 2004) and *dicp* genes (Haire et al. 2012). However, pure T and NK cell populations have so far not been isolated in zebrafish and no reliable antibody has been developed against orthologs of mammalian T and NK cell receptors. Therefore many properties of mammalian T and NK cell orthologs and their evolution in non-mammalian species remain uncharacterized.

High-throughput single cell RNA-Seq (scRNA-Seq) has emerged as a promising technology to unravel the landscape of cell types in heterogeneous cell populations without relying on specific antibodies (Saliba et al. 2014). Simultaneous expression of thousands of genes can be measured in each cell, thereby providing an unbiased view of transcriptional activity at the cellular level and avoiding the averaging effect of bulk gene expression studies (Shapiro et al. 2013). Cells can then be grouped into biologically relevant clusters based on similarity of their gene expression profiles rather than a handful of cell surface markers (Grün et al. 2015; Trapnell 2015; Macaulay et al. 2016). Therefore, despite technical and biological noise and the computational challenges associated with this variability (Brennecke et al. 2013; Buettner et al. 2015), scRNA-Seq has the potential to uncover new immune cell types that cannot be studied using traditional approaches.

To gain insight into the evolution of vertebrate immunity and non-mammalian immune cell types, we first analysed the conservation of mouse and human immune cell (i.e., T, B, NK and myeloid cells) specific genes. Next, we analysed immune cells in zebrafish, a powerful model in biomedical research (Langenau and Zon 2005; Renshaw and Trede 2012; Kaufman et al. 2016). To this end, we took advantage of a transgenic line of zebrafish expressing GFP under the control of the *lck* promoter (Langenau et al. 2004) and performed scRNA-Seq on *lck:GFP+* FACS sorted cells. Our analysis revealed three distinct *lck*+ cell populations: T cells, a novel type of NK-like cells and a smaller population of myeloid-like cells. Our expression profiles uncovered many immune signature genes both bony-fish specific and shared with mammals, including innate immune receptors, cytokines, transcription factors, proteases and antimicrobial peptides. In addition, we detected gene expression divergence among bony-fish duplicated paralogs. Finally, evolutionary analysis of differentially expressed genes showed higher gene turnover and lower conservation for NK/NK-like cells specific genes compared to T cells specific genes in all three species studied in this work.

## Results

### Conservation analysis of mammalian T, B, NK and myeloid cell specific genes across vertebrates

Immune related genes tend to evolve more rapidly than other genes and between functionally distinct immune cells the selective pressures might vary significantly. Here we performed a conservation analysis of the most differentially expressed genes in resting T, B, NK and myeloid cells in mouse and human at the genome-wide level (Chambers et al. 2007; Watkins et al. 2009) (see Methods).

Our analysis revealed that among trans-membrane (TM) or secreted protein coding genes, those specifically expressed in NK cells have proportionally less orthologs across all vertebrates compared to other immune cells. The difference is most evident between NK and T cells, although these are closer from a functional and ontogenical point of view (Fig. 1A and C). No difference, however, was observed for cytoplasmic or nuclear protein coding genes (Fig. 1B and D). As expected, the NK receptor families Ly49 in mouse and KIR in human strongly contributed to this difference. Interestingly, however, the differences between T and NK cell TM gene conservation were still observed after removing these receptors from the analysis (Supplemental Fig. S1). Examples of other mouse or human NK TM genes poorly conserved across vertebrates include Fc receptors, granulysin, CD160, CD244 and IFITM3. In addition, among conserved protein coding genes, NK cell specific genes consistently had lower sequence identity across all vertebrates for TM genes, but not for cytoplasmic ones (Supplemental Fig. S2).

**Fig. 1:**
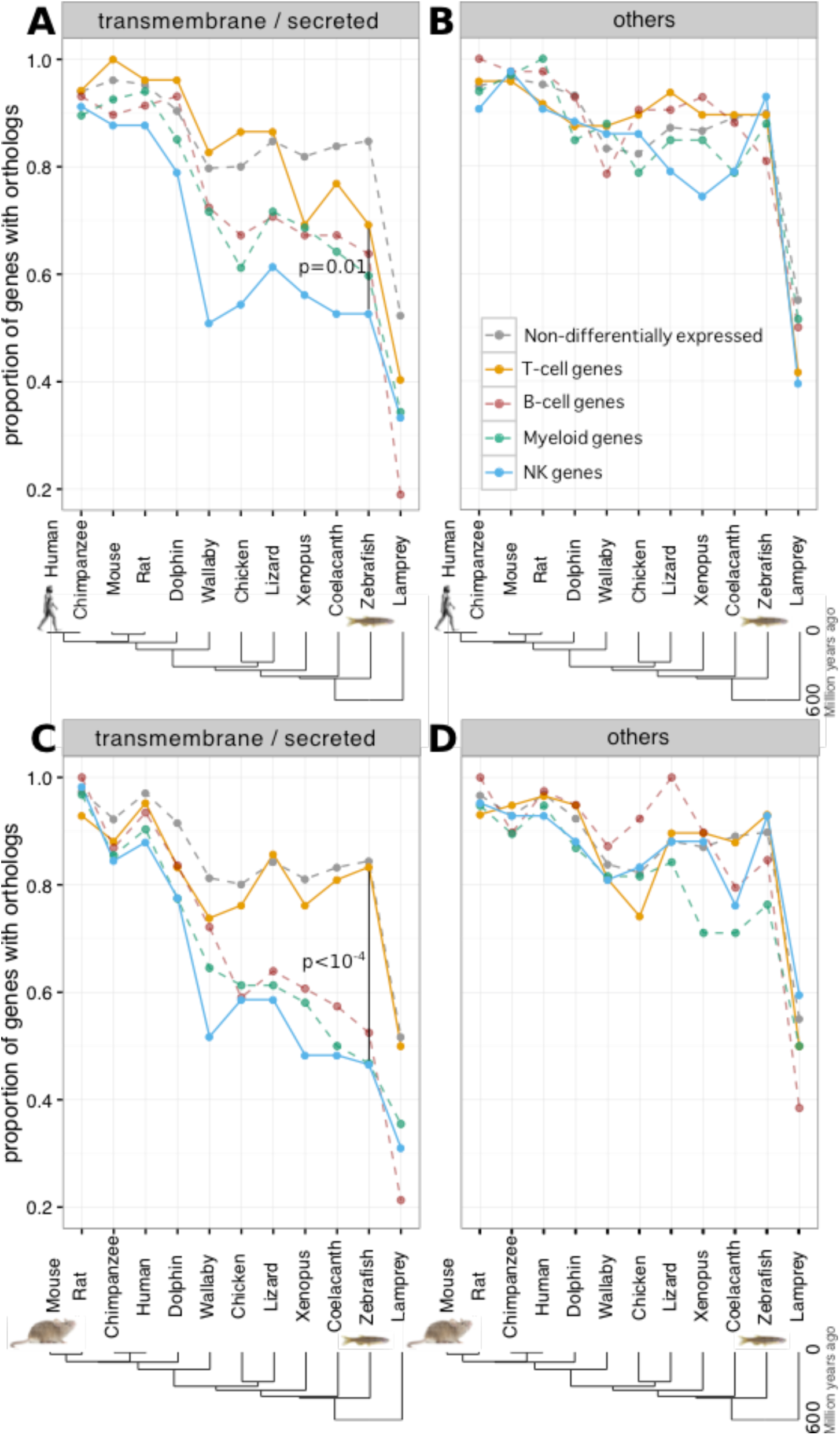
Conservation analysis of human and mouse genes differentially expressed in major immune cell types. A,B: Proportion of human genes specific for distinct immune cell types (T, B, NK and myeloid cells) with orthologs in other species. (A) shows the results for genes coding for transmembrane and secreted proteins and (B) for cytoplasmic and nuclear proteins. **C,D**: Same analysis as in (A) and (B) using mouse immune cell types’ specific genes.

The ratio between non-synonymous and synonymous substitutions (dN/dS ratio) of one-to-one orthologs between human and mouse, can provide a good estimation of the evolutionary pressure acting on a gene. Our results indicate that NKs’ TM genes evolve faster (i.e., present higher dN/dS values) compared to T cells’ TM genes (Supplemental Fig. S3). As expected, the lowest conservation for all immune cell type specific genes was observed in lamprey (Fig. 1) since these organisms possess a distinct adaptive immune system (Guo et al. 2009).

To further explore the conservation of immune cell types’ specific genes and expand our understanding of immune cell populations in an evolutionary distant non-mammalian species, we set out to profile immune cell populations in zebrafish.

### Single-cell transcriptomics of zebrafish lck+ lymphocytes reveals three distinct cell populations corresponding to T cells, NK-like cells and Myeloid-like cells

As reliable antibodies to isolate pure immune cell populations in fish species are not available, we used single cell transcriptome analysis of zebrafish *Tg(lck:GFP)* cells. This transgenic line expresses GFP under the control of the lymphocyte-specific protein tyrosine kinase (*lck*) promoter, and it was shown to be mainly restricted to zebrafish T cells (Langenau et al. 2004) However, as *Lck* in mouse and humans is expressed in both T and NK cells, we speculated that its expression pattern could be conserved in ray-finned fish. *Tg(lck:GFP)* zebrafish may therefore provide an ideal model to investigate the large difference in conservation between T and NK cell specific genes observed in mammalian species. To simultaneously obtain information about cell morphology and high quality gene expression profiles, we used high-throughput single-cell RNA sequencing combined with FACS (fluorescent activated cell sorting) index sorting analysis of two adult zebrafish (three and ten months old) spleen-derived *lck:GFP* cells.

We first generated and sequenced libraries from 278 single GFP+ cells isolated from the spleen of two different fish from a different clutch and different age (see Methods). Following quality controls (Supplemental Fig. S4, see Methods) 15 cells were removed and gene expression profiles for the remaining 263 cells were generated. Average single-cell profiles showed good correlation with independent bulk samples (PCC=0.82, Supplemental Fig. S5). Correlations between single-cell gene expression profiles were used to calculate cell-to-cell dissimilarities (see Methods) and these were represented into low dimensional space using classical Multidimensional Scaling (see Methods). Interestingly, a clear cell subpopulation structure emerged (Fig. 2A) showing three distinct cell groups. The three groups were confirmed by unsupervised whole-transcriptome clustering (see Methods, Fig. 3 and Supplemental Fig. S6E-F).

**Fig. 2:**
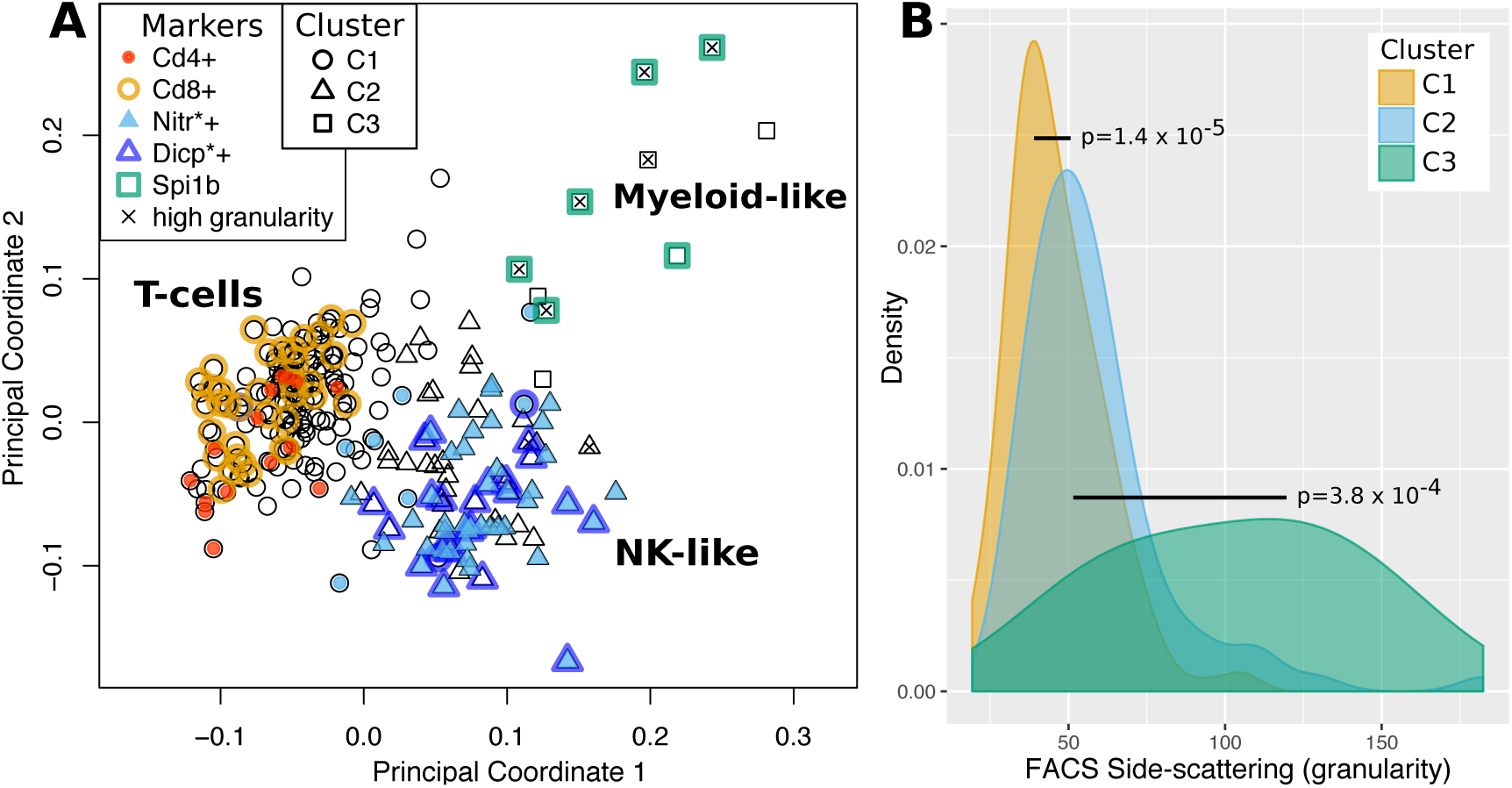
Multidimensional scaling of zebrafish *lck*+ single-cell transcriptomes. Unsupervised clustering revealed three subpopulations of cells: cluster 1 (C1), cluster 2 (C2) and cluster 3 (C3), containing 65%, 31% and 4% of the cells, respectively, and depicted with different symbols. A few examples of immune signature genes are depicted (using an expression threshold of 5 TPMs): *cd4* and *cd8* for T-cell specific genes, the innate immune receptors *nitr* and *dicp* for putative NK-like specific genes in zebrafish and the myeloid associated transcription factor *spi1b/pu.1* for myeloid-like cell specific genes. High granularity depicts cells with high side scattered light. **B:** Distribution of side scattered light (proxy for cellular granularity) for cells in each cluster.

**Fig. 3:**
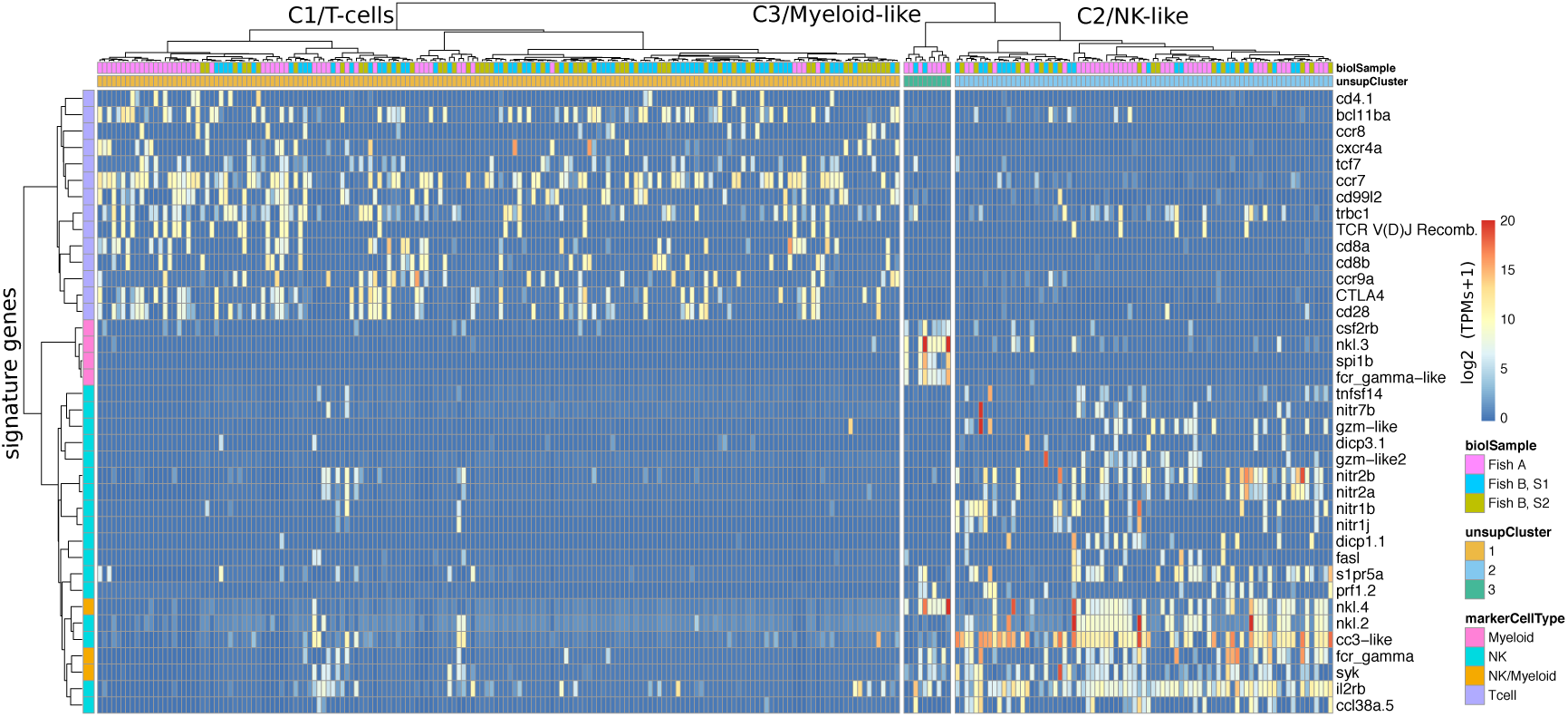
Heatmap showing the expression levels of some differentially expressed marker genes. Columns and rows represent cells and marker genes, respectively. Colours of the columns show the plates (top row) and the assigned clusters for each cell based on unsupervised whole-transcriptome clustering (second row, dendrogram shown on top). Colours of the rows (left-most column) indicate the known function of marker genes based on literature (T cell, NK or myeloid marker). The heatmap colour scale indicates the log2 transcripts per million (TPM, see Methods). Apart from a few cells in the T cell cluster that show expression of NK markers, the unsupervised whole-transcriptome clustering is very well recapitulated by expression of known and putative cell type markers.

The first cluster (C1) seemed to correspond to T cells based on the expression of *cd8a* and *cd4* genes (Fig. 2). To further support this hypothesis, we adapted a recent method for detection of V(D)J recombination events of the TCR locus (Stubbington et al. 2016) (see Methods). With a median of only 0.64 million gene-mapped reads per cell, we were able to unambiguously detect V(D)J recombination events in 27 cells (Fig. 3 and Supplemental Fig. S10). Occurrence of V(D)J recombination was associated with Cluster 1 (p<0.01, see Methods), which provides additional genomic evidences of the T cell identity. As expected, V(D)J recombined segments were also strongly associated with expression of the TCR beta chain constant region (*trbc1*) (p < 10^−5^, Fig. 3). Interestingly, *cd8* and *cd4* displayed mutually exclusive expression (as expected for mature T cells) (Fig. 2A and 3) and *cd4*+ and *cd8*+ cells clearly separated when low dimensional projection is restricted to cells from C1 (Supplemental Fig. S8).

The second cluster (C2) displayed expression of NK lysins, which have been recently proposed to mark a distinct sub-population of NK-like cells and are upregulated in Recombination activation gene 1 deficient (*rag1*−/−) zebrafish (Moore et al. 2016; Pereiro et al. 2015). In addition, multiple members of the teleost fish specific innate immune receptor families *nitr* (novel immune type receptors) and *dicp* (diverse immunoglobulin domain containing proteins) are highly expressed and specific to this cluster. It has been suggested that these receptors play a similar role as mammalian NK receptors (Yoder et al. 2004). Therefore, we hypothesized that this subpopulation corresponds to a zebrafish equivalent of mammalian NK cells.

Finally, a third cluster (C3) showed high expression of the myeloid lineage specific transcription factor (TF) *spi1b* (Ward et al. 2003). These data suggested that cells in Cluster 3 had a myeloid cell-like identity.

The clustering structure of our fish immune cells was further validated in a set of more than 300 single cells from a third fish and additional cells from the first fish, where despite much lower coverage due to external RNA contamination of the samples, the separation between cells expressing the different markers (*cd4*, *cd8*, *nitr*, *dicp* and *spi1b*) is clearly visible (Supplemental Fig. S7 and Supplemental Methods).

In addition to distinct transcriptional states, FACS analysis revealed that cells in different clusters differ in their light scattering properties (Fig. 2B). In particular, side scattered light (SSC), which is positively correlated with subcellular granularity or internal complexity, was 25% higher in C2 than in C1 (Wilcoxon test p = 1.4e-05). This is consistent with NK-like cells possessing dense cytoplasmic granules (Yoder and Litman 2011). In addition, SSC of cells in C3 was 203% higher than in the other two clusters together (p = 1.6e-05). The high granularity of cells in C3 further supports the hypothesis that these cells originate from a subpopulation of *lck+* myeloid cells, such as granulocytes (see (Gibbings and Befus 2009) for similar findings in mammals).

Since *lck:GFP+* cells were sorted randomly from spleen, the number of cells within each of the clusters could be used as an estimate of the frequency of each sub-population in the spleen in zebrafish. Similar to what is known from mouse (including *Lck:gfp* transgenic mice (Shimizu et al. 2001)) and human, T cells were more frequently found (65.4% of cells fall in C1) than NK-like cells (30.8% of cells fall in C2).

### Differential expression analysis identifies both known and novel genes specific for each cell type

To identify genes specific for each cell population we performed differential expression analysis of each cluster versus the other two (see Methods and Supplemental Table S1).

The T cell signature genes *cd4*, *cd8*a, *ctla4*, and cd28, the transcription factors *bcl11b* and *tcf7* and the cytokine/chemokine receptors *il10rb*, *ccr7*, *ccr9* and cxcr4 were within the most differentially expressed genes in Cluster 1 (Fig. 3). We also identified many T cell specific genes that were uncharacterized or did not have an informative name or description in the zebrafish genome for which we assigned a putative name, based on sequence similarity searches. These included the *cd8* beta chain (ENSDARG00000058682) whose expression is highly correlated with the alpha chain *cd8a* within Cluster 1, cd28 (ENSDARG00000069978) and an uncharacterized Ig-like protein (*ENSDARG00000098787*) related to CD7 antigen (Fig. 3).

Mammalian NK cells kill target cells by either of two alternative pathways: the perforin/granzyme secretory pathway or the death receptor pathway. Our analysis revealed differential expression of several members of both pathways in C2. For instance, differential expression of known innate immune receptors *nitr* and *dicp*, *syk* kinase, multiple granzymes, perforins and NK lysins is linked with activation of the secretory pathway whereas differential expression of FAS ligand (*faslg*) indicates activation of the death-receptor-ligand pathway (Dybkaer et al. 2007) in NK-like cells (Supplemental Table S1 and Fig. 3). Expression of these genes further shows that zebrafish presumably resting NK-like cells transcriptionally resemble effector CD8 T cells, as observed in mammals (Bezman et al. 2012).

We also observed a high expression level of cytokines and cytokine receptors. For example, differentially expressed genes in Cluster 2 included the sphingosine 1-phosphate receptor *s1p5* (*s1pr5a*, whose homolog in mammalian NK cells is required for homing), the interleukin-2 receptor beta (IL2 induces rapid activation of mammalian NK cells), *tnfsf14* (*Tnfsf14* tumor necrosis factor (ligand) superfamily, member 14) and chemokines of the families *ccr38* and *ccr34*. In addition, we detected differential expression of putative activating NK receptors’ adaptors (ITAMs) Fc receptor gamma subunit FcRγ (*fcer1g*) DAP10 (*hcst*), CD3ζ/cd247-like (*cd247l*) and multiple putative transcription factors (Supplemental Table S1). Finally, within the top differentially expressed genes of these NK-like cells we found putative homologs of mammalian granzyme B that is expanded in ray-finned fish genomes (ENSDARG00000078451, ENSDARG00000093990, ENSDARG00000055986, see also Fig. 4), and many uncharacterized putative Immunoglobulin-like receptors and cytokines, such as immunoglobulin V-set domain containing proteins or interleukin-8-like domain containing chemokines (Table 1). Altogether, these results add confidence in our proposed classification of these cells as putative fish NK-like cells.

**Fig. 4:**
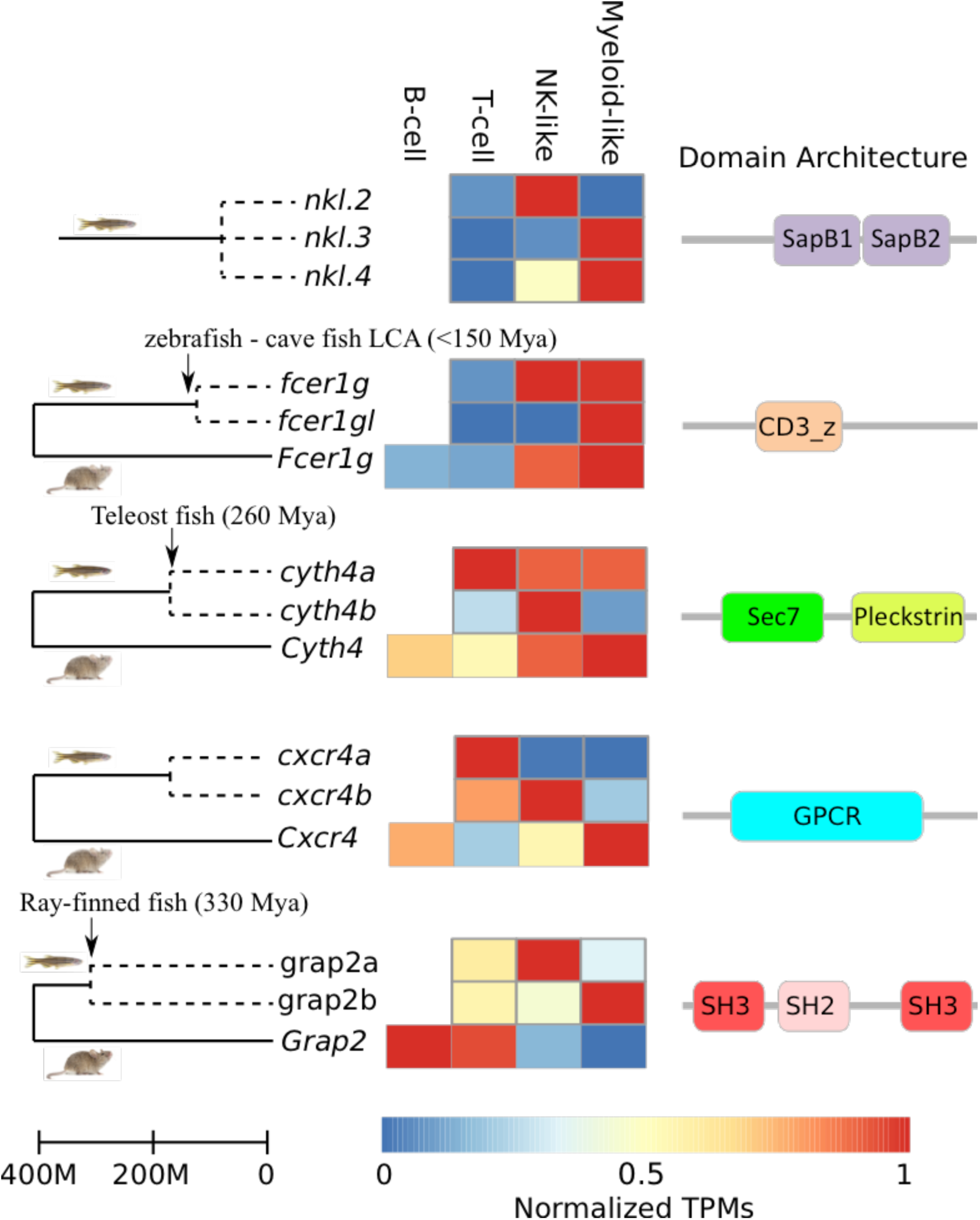
Examples of ray-finned fish-specific duplicated genes with diverged expression patterns. For genes with known mammalian orthologs, the expression in mouse is shown below. Times of gene duplication are indicated with arrows. Domain architectures were retrieved from PFAM (CD3_z: T cell surface glycoprotein CD3 zeta chain; GPCR: G-protein coupled receptor; SH2/3: Src-homology 2/3).

**Table 1.**
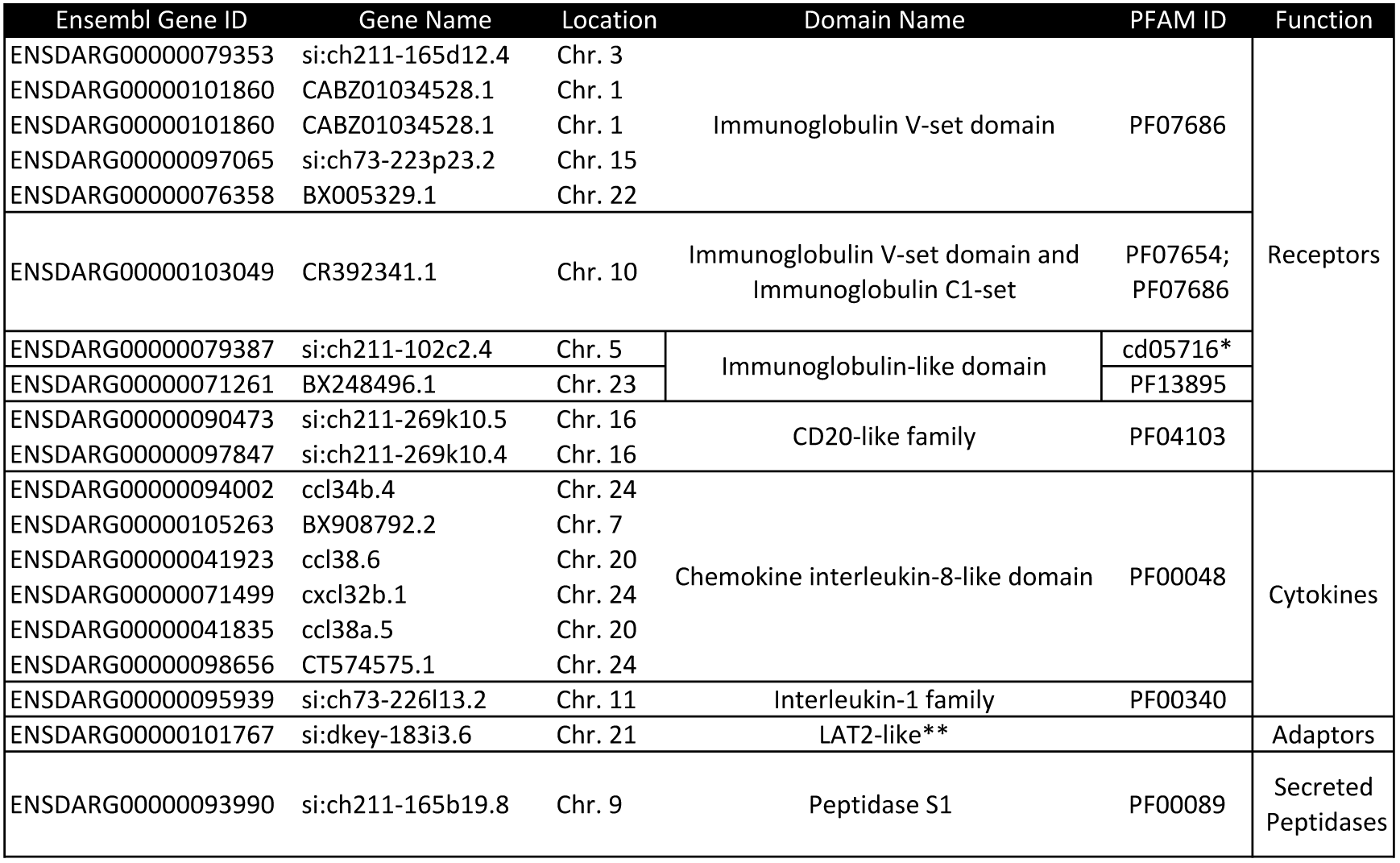
List of novel zebrafish NK-specific membrane-bound or potentially secreted proteins, including putative receptors, cytokines and related proteins. Domain annotations were retrieved from PFAM except for * (NCBI Conserved domains database) and ** (PSI-BLAST search).

Regarding cells in Cluster 3, the small number of cells within this cluster limits the power of differential expression analysis. Nevertheless, within the most differentially expressed genes in Cluster 3, we found two myeloid specific genes: the TF *spi1b* and the granulocyte/macrophage colony-stimulating factor receptor beta (*csf2rb*). Other differentially expressed genes included an Fc-receptor gamma like protein (*fcer1gl*), *hck*, a member of the Src family of tyrosine kinases mostly expressed by phagocytes in mammals and potentially implicated in signal transduction of Fc receptors and degranulation (Guiet et al. 2008), complement factor B (*zgc.158446*), and *id2* (a transcription factor interacting with *spi1b*), Fig. 3.

We next compared differentially expressed genes in each cell population to human transcriptomic data of homogeneous FACS sorted immune cells (Watkins et al. 2009; Chambers et al. 2007). For genes differentially expressed in Cluster 1, our results show a significant enrichment in differentially expressed in human T cells (P=0.008, see Methods). Similarly, the comparison of differentially expressed genes in Cluster 2 with human gene expression data confirmed a significant enrichment in NK specific genes (P=0.009) thus supporting the conservation of a core transcriptional program between mammalian and zebrafish NK-like cells (see Methods). Finally, differentially expressed genes in Cluster 3 were weakly enriched in human Myeloid-specific genes (odds ratio=5.2, P=0.06, see Methods).

### Functional divergence of duplicated immune genes in zebrafish

Gene duplication is a common event in eukaryotic genomes and plays a major role in functional divergence. To systematically explore this functional divergence in fish immune genes, we collected all duplicated genes pre- and post-ray finned fish speciation (see Methods). Interestingly, genes more recently duplicated (ray finned fish-specific) show lower expression in our dataset. For example, 53% of pre-speciation duplicated genes showed expression in *lck+* cells, compared to 41% of post-speciation duplicated paralogs. As expected, pre-speciation duplicated immune genes were more likely (94%) to functionally diverge (i.e. show differential expression in the immune subpopulations, see Methods) compared to the more recent post-speciation paralogs (62%). Ray finned fish-specific duplicated genes with conserved expression patterns included, for instance, the NK receptors *nitr* that, although expanded in zebrafish, have kept their cell type specificity. In contrast, other fish-specific paralogs show distinct expression, suggesting possible neo-functionalization events (see Fig. 4). NK-lysins (*nkl.2*, *nkl.3*, *nkl.4*) provide an interesting example of recent functional divergence. In our data *nkl.4* was expressed in both Myeloid- and NK-like cells. However, *nkl.3* was only expressed in Myeloid-like cells, while *nkl.2* expression was restricted to NK-like cells (Fig. 3 and 4). A second example of neofunctionalization is the Fc receptor gamma subunit (FcRγ), which in mouse and human, is highly expressed in myeloid and NK cells (Tassi et al. 2006). In zebrafish *lck+* cells, fcr_gamma (*fcer1g*) was expressed in Myeloid- and NK-like cells, while its paralog fcr_gamma-like (*fcer1gl*) expression was restricted to the Myeloid-like cells (Fig. 4). Other examples of such neo- or sub-functionalization of recently duplicated paralogs are shown in Fig. 4 and Supplemental Table S2.

### NK specific genes show lower conservation than T cell genes from mammals to bony fish

The immune system is constantly adapting to new pathogens and changes in virulence mechanisms, and hence is one of the most rapidly evolving biological systems in vertebrates (Fumagalli et al. 2011; Kosiol et al. 2008). To explore the evolution of the newly identified zebrafish genes specific for T, NK-like and myeloid-like cells, we performed the same conservation analysis as in Fig. 1 (see Methods). Consistently, among TM or secreted proteins, 76% of differentially expressed genes in zebrafish T cells had orthologs in mouse or human compared to ~36% of differentially expressed genes in zebrafish NK-like cells (p<10^−4^, Fig. 5), suggesting higher rates of gene turn-over in NKs across vertebrate evolution. Among non-TM or secreted protein coding genes, proportion of orthologs was similar between T, NK-like and myeloid-like cell specific genes (Fig. 5), as observed in mammalian species.

**Fig. 5:**
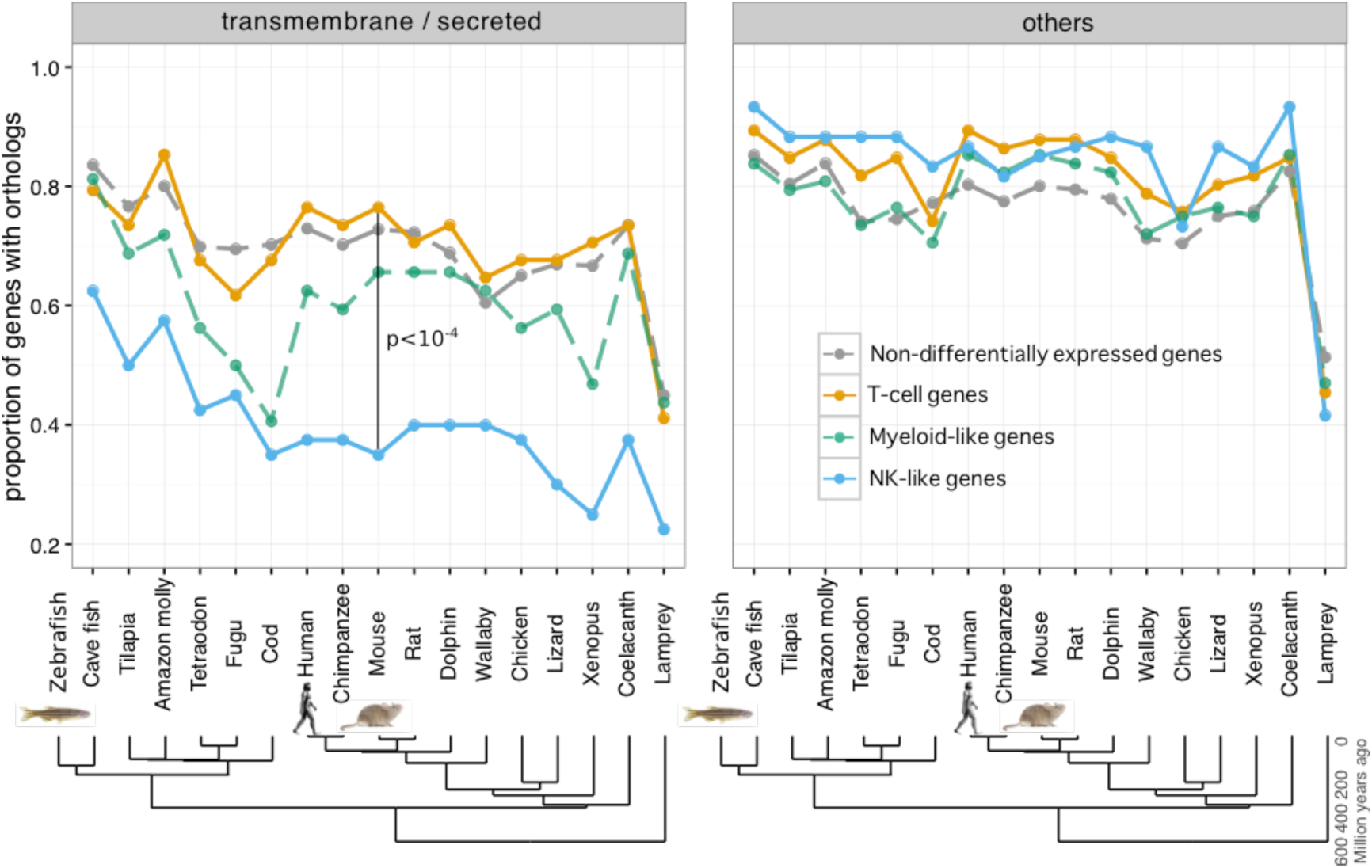
Conservation analysis of zebrafish immune genes across vertebrates. The proportion of orthologs of protein-coding genes for non-differentially expressed genes (grey), differentially expressed genes in T cells (orange), NK-like cells (blue) and myeloid-like cells (green) are shown for both transmembrane or secreted proteins (left) and other proteins (right).

Examples of TM genes with no assigned orthologs beyond bony fish include putative chemokine receptors (e.g. ENSDARG00000105363), nk lysins and the NK receptors *nitr* and *dicp* among NK-like specific genes (see also Table 1), as well as the Ig-like protein coding genes (e.g. ENSDARG00000098787, ENSDARG00000092106 and ENSDARG00000092106, Supp. Table 1) among T cell specific genes. Although *lck*+ myeloid cells represent only a sub-population of fish myeloid cells, their genes consistently show intermediate level of conservation between T and NK cell specific genes, as observed for myeloid cells in mammals (Fig. 1 and 5).

When compared at the sequence identity level, the conserved TM genes specifically expressed in either T or NK-like cells had lower sequence identity than other genes across vertebrates (p<10^−6^, Supplemental Fig. S2C). Moreover, as in mammals, zebrafish TM genes were on average less conserved in NK-like than T cells across vertebrates (p=0.03, Supplemental Fig. S2C). In contrast, cytoplasmic and nuclear T cell specific displayed similar sequence identity compared to other genes (Supplemental Fig. S2C)

## Discussion

The availability of fully sequenced genomes in several vertebrate species has enabled analysis of the evolution of the immune system based on orthology of known mammalian immune cell markers. However, a comparison of immune subtypes at the cellular and molecular level has progressed slowly, mainly owing to the lack of suitable antibodies that mark distinct immune cell subpopulations in lower vertebrates. Here we used scRNA-Seq of *lck:GFP* cells to characterise immune cell subpopulations in zebrafish and examine their conservation in other vertebrate species. Our work establishes scRNA-Seq as a powerful technique to study immune cell types across vertebrate species.

Using single cell from two fish, we find three consistent clusters of cells, each comprising cells from both fish. The most abundant population of cells in our data set had a clear molecular T cell signature. The cells in this cluster showed differential expression of hallmark genes important in regulation of T cell development and signalling, suggesting a conserved transcriptional program from mammals to fish. Within this population we were able to detect TCR V(D)J recombination in 22 cells (Fig. 3 and Supplemental Figure 10). Interestingly, a single TCR recombinant was found in each cell (Supplemental Table 5), which is consistent with allelic exclusion. Although V(D)J recombination was clearly correlated with T cell identity, five cells with evidence of V(D)J recombination fall in Cluster 2 and three of them show clear expression NK genes. It is tempting to speculate that these cells could be NKT cells. However, in mammals, the process of TCR rearrangement first initiates in uncommitted haematopoietic progenitors before NK/DC/B/T divergence. Therefore, incomplete rearrangements are also observed in subpopulations of non T cells, such as NKs (Pilbeam et al. 2008). This could explain the presence of V(D)J rearrangements in NK-like cells at the transcriptional level, as well as expression of single V or J segments in cells in Cluster 2 and Cluster 3. Moreover, TCRs expressed by NK T cells present a limited diversity while here we found no evidence for preferential use of specific segments among these cells.

In mammals LCK is expressed in both T and NK cells and in our dataset one population of *lck+* cells resembled NK cells. Although NK-like cells were first identified in catfish (Shen et al. 2004) over a decade ago, very little is known about the NK cell transcriptome beyond mammals. Our data revealed that the proposed bony fish NK receptors of the *nitr* family showed restricted expression in a distinct cell subpopulation of NK-like cells which also expressed granzymes, perforins, NK lysins (Pereiro et al. 2015; Moore et al. 2016), FAS ligand, TNFSF14, IL2 receptor beta, the homolog of chemokine receptor CCR2, the sphingosine 1-phosphate receptor S1P5 (required for homing of mammalian NK cells), specific transcription factors and multiple novel putative NK-specific receptors and chemokines (Fig. 3 and Table 1).

Throughout evolution, animals and plants have developed complex immune defence mechanisms to combat microbial infections. However, pathogens experience strong selective pressure to evade host recognition and thus impose selective pressure on the host to re-establish immunity. As a consequence, immune-related genes have been preferential targets of positive selection in vertebrates (Kosiol et al. 2008; Yoder and Litman 2011). Using a genome-wide unbiased approach based on transcriptomic data from two mammalian and one bony fish species, we showed that a lower fraction of orthologs and lower protein sequence identity are observed for NK TM genes compared to other immune cell type specific TM genes, and especially T cell TM genes, even though T and NK cells are functionally more related (e.g., TCD8 and NK cytotoxicity upon MHCI recognition). Importantly, the trend is not only due to known NK receptors (i.e. Ly49 in mouse, KIR in humans and *nitr/dicp’*s in zebrafish, Supplemental Fig. S1). This suggests that rapid evolution of NK TM genes is key for their function in all vertebrates. As NK genes cannot undergo somatic rearrangement, we propose that this fast evolution reflects, at least partly, a need for NK cells to possess a diverse repertoire of species-specific germline encoded receptors and associated proteins to perform their functions. In particular, both T and NK cells recognize the fast evolving and highly polymorphic MHC molecules. While T cells do so by rearranging their TCR sequence, NK cells possess an expanded family of receptors. The fast evolution of these receptors may be the result of a need to adapt to MHC rapid evolutionary changes. Our observations also support a model of high gene turnover and faster evolution of immune TM/secreted genes, but at the same time conservation of core cytoplasmic immune genes from zebrafish to mammalian species (Fig. 1 and 5). As such, it supports zebrafish as an appropriate model organism for immune cell intracellular signalling studies.

Overall, our work expands the analysis of immune subpopulations and their evolution to lower vertebrates. To our knowledge, this is the first study to characterize T and NK cells at the transcriptome level in a non-mammalian species and one of the first studies to analyse NK cells’ gene expression at the single cell level (see (Moore et al. 2016) for qPCR analysis of *TG(lck:GFP)* zebrafish single cells). We confirmed cell-type specific expression of expected zebrafish T and NK cell genes and predicted new markers of these two cell types. We further found significant variability in highly expressed NK receptors and identified multiple cases of immune genes duplications followed by neofunctionalization. Global conservation analysis revealed more rapid turnover of NK specific TM genes compared to other immune cell, and especially T cell specific genes in mammals and fish suggesting that this is an essential property of immune cells.

## Methods

### Conservation analysis of mouse and human immune cell differentially expressed genes

Orthologs of mouse and human protein-coding genes and their sequence identities, as well as transmembrane domains and signal peptide predictions were downloaded from BioMart / Ensembl Genes 82. For genes having multiple orthologs, we considered their average sequence identity. Mouse and human NK, T cell, B-cell, granulocyte and monocyte microarray gene expression datasets were obtained from (Chambers et al. 2007) and (Watkins et al. 2009). First we pre-filtered genes with low expression levels among these cell types using a threshold on normalized expression level of 5 for the mouse data (16060 genes), and 8 for the human data (8242 genes). CD8 and CD4 T-cells samples were merged into a T-cell group and Monocyte and Granulocyte samples were merged into a myeloid cells group. We then obtained differentially expressed genes in each group compared to the others, using limma (version 3.28.14). Significantly differentially expressed genes (Benjamini-Hochberg adjusted p-value < 0.01) were ordered based on expression fold-change and the top 100 genes unique for each cell type were selected as ‘signature genes’ for downstream analysis (Supplemental Table S4, sheets 2 and 3). Results were robust to different cut-offs for the top N differentially expressed (Supplemental Methods and Supplemental Fig. S9). Human and mouse dN/dS ratios (Supplemental Fig. S3) of one-to-one orthologs between these two species were obtained from Ensembl version 82. The two protein groups enriched in 1) transmembrane and secreted proteins and 2) cytoplasmic and nuclear proteins were defined based on the presence of predicted trans-membrane domains and/or signal peptide. Statistical significances of differences in sequence identity and dN/dS differences were assessed using Wilcoxon/Mann-Whitney tests. Statistical significances of differences in proportion of orthologs in 1) a specific species (e.g. ‘human’ point in Figure S2A) were assessed by comparison against a null-model distribution generated from 10,000 random permutations of gene – cell type specificity class pairs, and 2) globally across all species (as in Fig. S2 C), using paired Wilcoxon signed rank test (to evaluate ‘consistency’ of the difference in conservation patterns between two cell types).

### Zebrafish strains and maintenance

Wild type (Tubingen Long Fin) and transgenic zebrafish *Tg(lck:EGFP)* lines were maintained as previously described (Bielczyk-Maczyńska et al. 2014), in accordance with EU regulations on laboratory animals.

### Single cell sorting and whole transcriptome amplification

The spleens from two heterozygote *Tg(lck:EGFP)* adult fish from a different clutch and different age (3 and 10 months old) and one adult wild-type fish were dissected and carefully passed through a 40µm cell strainer using the plunger of a 1-mL syringe and cells were collected in cold 1xPBS/5% FBS. The non-transgenic line was used to set up the gating and exclude autofluorescent cells. Propidium iodide (PI) staining was used to exclude dead cells. Individual cells were sorted, using a Becton Dickinson Influx sorter with 488- and 561 nm lasers(Schulte et al. 2015) and collected in single wells of 96 well plates containing 2.3 uL of 0.2 % Triton X-100 supplemented with 1 U/uL SUPERase In RNAse inhibitor (Ambion). The size, granularity and level of fluorescence for each cell were simultaneously recorded. Seven wells were filled with 50 cells each, from the second fish to compare single-cell with bulk RNA-Seq (Supplemental Figure S5). The Smart-seq2 protocol (Picelli et al. 2014) was used to amplify the whole transcriptome and prepare libraries. Twenty-five cycles of PCR amplification were performed. Similar analysis was performed on two additional plates of the first fish and four plates from a third fish, including 5 wells with 50 cells (see Supplemental Methods and Supplemental Fig. S7).

### Single cell RNA-Seq data processing

Following Illumina HiSeq2000 sequencing (125bp paired-end reads), single-cell RNA-Seq reads were quality trimmed and cleaned from Nextera adaptor contaminant sequences using BBduck (*http://sourceforge.net/projects/bbmap*) with parameters *minlen=25 qtrim=rl trimq=10 ktrim=r k=25 mink=11 hdist=1 tbo*.

An average of 2.1 million paired-end reads were obtained per single-cell (Supplemental Fig. 4 B). Next, gene expression levels were quantified as E_*i,j*_=log_2_(TPM_*i,j*_+1), where TPM_i,j_ refers to transcript-per-million (TPM) for gene *i* in sample *j*, as calculated by RSEM 1.2.19 (Li and Dewey 2011). RSEM (which uses Bowtie 2.2.4 for alignment) was run in paired-end non strand-specific mode with other parameters by default using the latest zebrafish genome assembly and transcript annotations (GRCz10 / GCA_000002035.3) combined with eGFP sequence appended as an artificial chromosome. For each single-cell, ~0.8 million reads on average (with a median of 0.65 million) were mapped to the transcriptome (Supplemental Fig. S3A). On average, 1240 expressed genes per cell were detected (Supplemental Fig. S3C). Cells having less than 500 detected genes or less than 10,000 reads mapped to transcripts were excluded from further analyses.

### Transcriptome dimensionality reduction, batch effect removal and cell clustering

In order to visualize cell heterogeneity at the transcriptomic level, we used classical multidimensional scaling (MDS, *aka* Principal Coordinates Analysis; as implemented in R’s *cmdscale* function) for dimensionality reduction (Fig. 2A, Supplemental Fig. 6A). MDS attempts to preserve distances between points generated from any dissimilarity measure. Pearson’s correlation coefficients (PCC) between full transcriptional profiles were used to define cell-to-cell similarities, and 1-PCC was then used as MDS’s input dissimilarity measure. Similar low-dimensionality projection was obtained with Principal Components Analysis (PCA) on the expression levels (E_*i,j*_) of the 1500 most variable genes (Fig. 6B).

To correct for batch effects and remove unwanted variation between the first and second fish, we used ComBat function from R Bioconductor’s *sva* package (Parker et al. 2014). After this procedure, variation between individuals was minimal (Supp. Fig. 6G).

To identify different cell populations, we performed hierarchical clustering using Ward’s criteria (as implemented in R’s *hclust* using *Ward.D2* method) applied on the first four Principal Coordinates generated by the MDS. The choice of the components was based on the eigenvalue decomposition of the MDS (Supp. Fig. 6C). Eigenvalues decrease smoothly after the fourth Component, *i.e.* contributing less significantly to the overall variability. The three cell clusters C1, C2 and C3 (Fig. 2A, Fig6 E and F) were obtained by maximising the mean silhouette coefficient for different number K of clusters (Supp. Fig. 6D).

### TCR reconstruction

All four TCR loci (α, β, δ, and γ) and Rag-dependent variable diversity joining V(D)J recombination are found in zebrafish (Langenau and Zon 2005). However, only the beta chain locus was fully annotated (Meeker et al. 2010). To explore TCR recombination in our immune cell subpopulations, we adapted the recent method of (Stubbington et al. 2016). Synthetic beta chain sequences containing all possible combinations the 52 V and 33 J germline segments were generated, with the addition of 20 ‘N’ ambiguity bases in the 5’ end, 7 ‘N’s between V and J segments and 50 ‘N’s at the 3’ end to account for unknown leader, possible D and constant sequences, respectively. RNA-seq reads from each cell were aligned against the collection of synthetic TCR beta chain sequences independently using the Bowtie 2 aligner, with low penalties for introducing gaps into either the read or the reference sequence or for aligning against N nucleotides (parameters ‘--no-unal -k 1 --np 0 --rdg 1,1 --rfg 1,1’). Next, reads aligning to synthetic sequences were used as input to the Trinity RNA-seq assembly software (Grabherr et al. 2011) using its default parameters for *de novo* assembly. Contigs assembled by Trinity were used as input to NCBI IgBlast 1.4 (Ye et al. 2013) using parameters ‘-qcov_hsp_perc 90 -evalue 0.001 -ig_seqtype TCR -perc_identity 99‘ and providing zebrafish V, D and J segments, and the resulting output were processed with a custom parsing script. Contigs with no stop codons and matching both a V and a J segment with at least 90% sequence identity against corresponding germline segments, and where at least 90% of the germline segment was recovered, were considered evidence for TCR beta chain V(D)J recombination.

### Differential expression analysis and marker gene discovery

Estimated gene counts obtained from RSEM were used as input for *SCDE* R package v1.99 (Kharchenko et al. 2014) that explicitly accounts for high rate of dropout events in scRNA-Seq. Differential expression between each cluster versus the other two was assessed using 100 randomizations (Supplemental Table S1).

To assess transcriptional conservation between mammalian and zebrafish immune cell types, we used the previously defined sets of human top 100 differentially expressed genes in T, NK and myeloid cells. We then compared the proportion of zebrafish genes with orthologs in T cell, NK cell and myeloid signature genes within the differentially expressed genes in each cluster versus non-differentially expressed genes. Statistical significance was assessed using Fisher’s exact test.

### Expression analysis of duplicated immune genes in zebrafish

A list of paralogs in zebrafish was obtained from Ensembl Compara GeneTrees (Vilella et al. 2009) (version 82). We defined two groups of protein coding genes: 1) 14,342 genes that underwent ‘recent’ duplication, whose most recent common ancestor was mapped to ray-finned fish (Actinopterygii) or any of its child nodes (Neopterygii, Otophysa, Clupeocephala, *Danio rerio*), and 2) 19,499 genes that underwent ‘early’ duplication, where their most recent common ancestor was mapped to bony vertebrates (Euteleostomi) or any of its parent taxa (Bilateria, Chordata, Vertebrata). Many of these genes suffered multiple duplication events both before and after the fish common ancestor. Therefore, to compare differences in expression between these two groups, we did not include the set of overlapped genes and obtained 3,235 unique recently duplicated genes and 8,609 unique early duplicated genes. From these, 1315 (41%) and 4,569 (53%) were detected in our data (genes with > 0 TPM in at least 1% of the cells).

For the analysis of expression pattern divergence, we searched pairs of paralogs where both genes show some specific expression pattern (therefore, likely to have an immune-related function) according to one of the following criteria: 1) within the top 100 differentially expressed genes in Cluster 1, Cluster 2, or Cluster 3; 2) within the top 100 differentially expressed genes in Cluster 2 and Cluster 3 versus Cluster 1 (*i.e.*, depleted in Cluster 1), Cluster 1 and Cluster 3 versus Cluster 2 (*i.e.*, depleted in Cluster 2), or Cluster 1 and Cluster 2 versus Cluster 3 (*i.e.*, depleted in Cluster 3); 3) expressed in all the three clusters (in at least 10% of the cells of each cluster). In the latter case, we only considered pairs of paralogs where only one gene is expressed in the 3 clusters, and the second is either specifically expressed or depleted from the major clusters 1 or 2 (pairs of paralogs where both genes are expressed in all 3 clusters were not considered since most of them are not immune-related genes, and cluster 3 is too small to accurately assess enrichment/depletion).

Next, we identified cases where both paralogs belong to the same expression pattern group (duplicate genes with conserved expression pattern) and cases where they differ (cases of neofunctionalization due to different expression patterns). For recently duplicated genes we found 23 pairs with distinct expression patterns and 14 pairs with the same expression patterns (i.e. 62% of paralogs’ neofunctionalization), while for early duplicated genes we found 121 pairs with distinct patterns and 8 pairs with the same expression patterns (i.e. 94% of paralogs’ neofunctionalization), as shown in Supplemental Table S2.

### Gene sequence conservation analysis of zebrafish differentially expressed genes

Orthologous genes of zebrafish in vertebrate species and their sequence identities were downloaded from BioMart / Ensembl Genes 82. For comparisons between differentially expressed genes between Cluster 1 (T cells), Cluster 2 (NK-like cells) and Cluster 3 (myeloid-like cells) we chose the top 100 differentially expressed genes after filtering by Z-score > 1 and sorting by fold-change (Supplemental Table S1). Results were robust to different cut-offs (Supplemental Methods and Supplemental Fig. S10). To assess ortholog conservation among non differentially expressed genes, we first excluded lowly expressed genes from the analysis (those where its expression level E was below the global mean of 0.46). The reason for this is that we observed a bias of higher gene conservation among highly expressed genes compared to lowly expressed genes. After this filter, conservation of differentially expressed genes could be compared to that of non-differentially (but having equivalent expression levels) genes as in Figure 5.

### Data access

The data reported in this paper was deposited in ArrayExpress under the accession number E-MTAB-4617

## Acknowledgements

With thank Prof. Pedro Romero for a critical reading of the manuscript.

The study was supported by SystemsX (MelanomX grant for S.C.), Cancer Research UK grant number C45041/A14953 to A.C. and L.F, European Research Council project 677501 – ZF_Blood to A.C. and a core support grant from the Wellcome Trust and MRC to the Wellcome Trust – Medical Research Council Cambridge Stem Cell Institute. Computations were performed at the Vital-IT (http://www.vital-it.ch) Center for high-performance computing of the Swiss Institute of Bioinformatics.

